# Short-term consumption of sucralose with, but not without, carbohydrate impairs neural and metabolic sensitivity to sugar

**DOI:** 10.1101/557801

**Authors:** Jelle R. Dalenberg, Barkha P. Patel, Raphael Denis, Maria G. Veldhuizen, Yuko Nakamura, Petra C. Vinke, Serge Luquet, Dana M. Small

## Abstract

There is a general consensus that overconsumption of sugar sweetened beverages contributes to the prevalence of obesity and related comorbidities such as type 2 diabetes (T2D). Whether a similar relationship exists for no, or low-calorie “diet” drinks is a subject of intensive debate and controversy. Here, we show that metabolic dysfunction, coupled with reduced central sensitivity to sweet, but not sour, salty or bitter taste, occurs when sucralose is repeatedly consumed with, but not without, a carbohydrate over a two-week period in healthy humans. A similar exposure to sucralose, with, but not without, a carbohydrate altered substrate utilization in mice. More specifically, greater energy intake was required for the animals to shift from fatty acid to carbohydrate oxidation, indicating a reduced sensitivity to carbohydrate. These findings demonstrate that consumption of sucralose in the presence of a carbohydrate rapidly impairs glucose metabolism and may contribute to the rise in T2D.

## Introduction

Significant controversy exists over the effects of consuming no, or low-calorie sweeteners (LCS) on health. Human studies have reported that consumption of LCS is positively associated with weight gain and/or diabetes [1–4], positively associated with lower BMI and weight loss [5–7], or unrelated to metabolic and body weight measures [8,9], possibly due to methodological limitations [10]. A similar inconsistency exists in the animal literature, with three recent reviews reaching three different and mutually exclusive conclusions [1,9,11]. Given the growing use of LC [12], especially in relation to the obesity and diabetes pandemics, it is of pressing importance to resolve the controversy surrounding LCS consumption.

Central to resolving this debate is defining and testing biologically plausible mechanisms by which LCS could lead to metabolic impairment. Several have been proposed [13–16]. The binding of LCS to extra-oral taste receptors in the pancreas and intestine could influence glucose absorption by affecting glucose transporters SGLT-1 and GLUT2 or by altering glucose metabolism by promoting incretin release. Central mechanisms could also play a role. For example, it has been suggested that uncoupling sweet taste from energy receipt leads to a weakening of conditioned responses to sweet taste [17]. In this case, sweetness-elicited conditioned responses, such as release of incretins, which help regulate glucose metabolism, is hypothesized to be reduced, leading to the subsequent development of glucose intolerance [17]. Support for this uncoupling hypothesis comes from a series of studies in rodents reporting weight gain or glucose intolerance in rats consuming yogurts sweetened inconsistently with sucrose and LCS compared to rats consuming yogurts consistently sweetened with only sucrose [14,18–22].

In the current study we set out to test the sweet uncoupling hypothesis in humans. Forty-five healthy humans were randomly assigned to consume: (1) beverages sweetened with sucralose (sweet uncoupled from calories - LCS), (2) beverages sweetened with maltodextrin (sweet coupled with calories - Carb), or (3) beverages sweetened with sucralose and combined with maltodextrin (Combo). Oral glucose tolerance test (OGTTs), sensory tests, and neuroimaging were conducted before and after participants consumed seven of their assigned beverages over 2-weeks in the laboratory. We reasoned that if the uncoupling hypothesis is correct, then participants in the LCS, but not the Carb or Combo groups should have reduced insulin sensitivity coupled with decreased brain and sensory response to sweet, but not sour, salty or savory taste. A parallel study was conducted in adolescents and a follow-up study in mice.

## Results

### Human Studies

Forty-five healthy young adults aged 18-45 who were non-regular consumers of LCS. A parallel study was conducted with adolescents, aged 13-17, since adolescents go through a period of transient insulin resistance [23], a time of increased preference for sweet beverages and of intensive brain development [24–28], especially for dopaminergic and prefrontal cortical circuits [29]. In these studies, we assessed glucose tolerance and taste perception before and after participants consumed seven 355ml novel-flavored equi-sweet beverages over two weeks using randomized double-blind designs. These beverages were sweetened with either 0.06g sucralose (0 Kcal, uncoupled stimulus), equi-sweet 30.38g sucrose (120 Kcal, coupled stimulus) or a control beverage containing the same dose of sucralose plus 31.83g of the non-sweet carbohydrate maltodextrin (120 Kcal, coupled stimulus). In addition, we measured brain response to sweet, sour, salty and savory taste using functional magnetic resonance imaging (fMRI). We reasoned that if uncoupling sweet taste from energy affects sweet taste guided feeding and conditioned responses, then uncoupling should result in glucose intolerance and reduced brain and perceptual responses to the sweet taste of sugar relative to other tastes in the uncoupled stimulus group, but not in the other two groups. A study overview is given in **Figure 1**. Detailed participant demographics are provided in **Table S1.**

**Figure 1:**
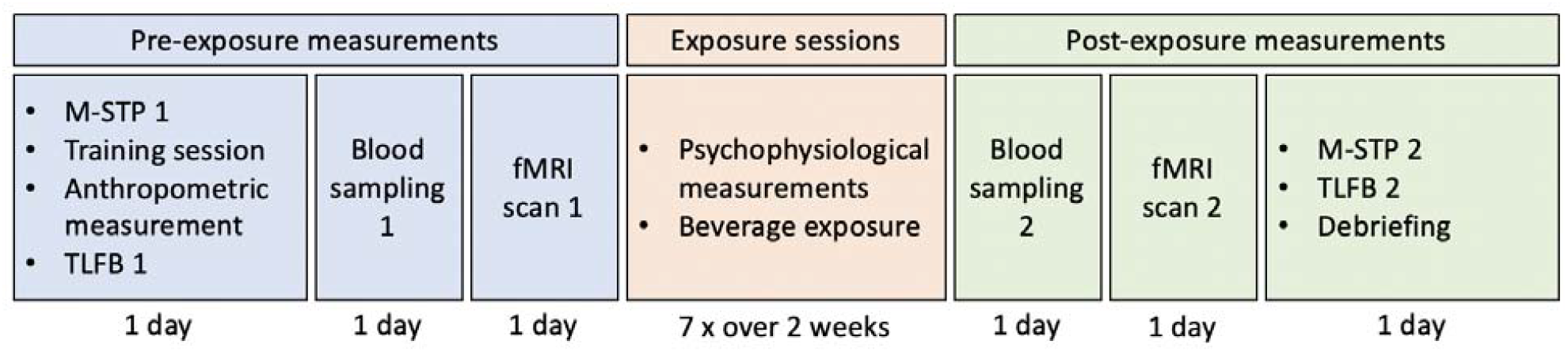
Human Study overview. Participants visited the lab 13 times. Measurements were divided into pre-exposure measurements, exposure sessions and post-exposure measurements. NQ: Nutrition Questionnaire; M-STP: Monell forced-choice sweet taste preference test; TLFB: time line follow back; fMRI: functional magnetic resonance imaging.

### Insulin sensitivity is reduced following consumption of sucralose with, but not without, saltodextrin

Glucose tolerance was assessed in young adults using the incremental area under the curve (iAUC) of blood plasma insulin and glucose during an Oral Glucose Tolerance Test (OGTT). We found a significant difference between the groups for first phase insulin response (time 0-30 min, F(2,36)=3.88, P=0.03) (**Figure 1A**) while we found no group differences in the first phase glucose response (F(2,36)=0.43,p=0.65) (Figure S1 and Stars Methods). Contrary to the uncoupling hypothesis, post hoc tests revealed a larger first phase insulin response in the Combo group (i.e., exposed to sucralose plus maltodextrin) compared to the LCS and Carb groups (exposed to sucrose alone or sucralose alone; false discovery rate corrected *t* tests; β=37.00%, P=0.03 and β=39.59%, P=0.03, respectively). In the adolescent group, glucose tolerance was assessed using a single timepoint blood draw to measure fasting blood plasma insulin and glucose. Based on our findings in the adults we halted the adolescent trial to examine the data for adverse effects. We found that HOMA-IR levels elevated from <3.5 to >12.9 in 2 out of 3 participants in the sucralose plus maltodextrin group. This elevation was driven by an increase in fasting blood plasma insulin levels (**Figure 2B**). We reported this adverse event to the Human Investigations Committee, which recommended trial termination. While the small group numbers currently do not permit us to draw any firm conclusions, permutation testing (n=1000) indicated that the HOMA-IR difference scores of the Combo group are significantly different from the LCS and Sugar groups together (p=0.043).

**Figure 2.**
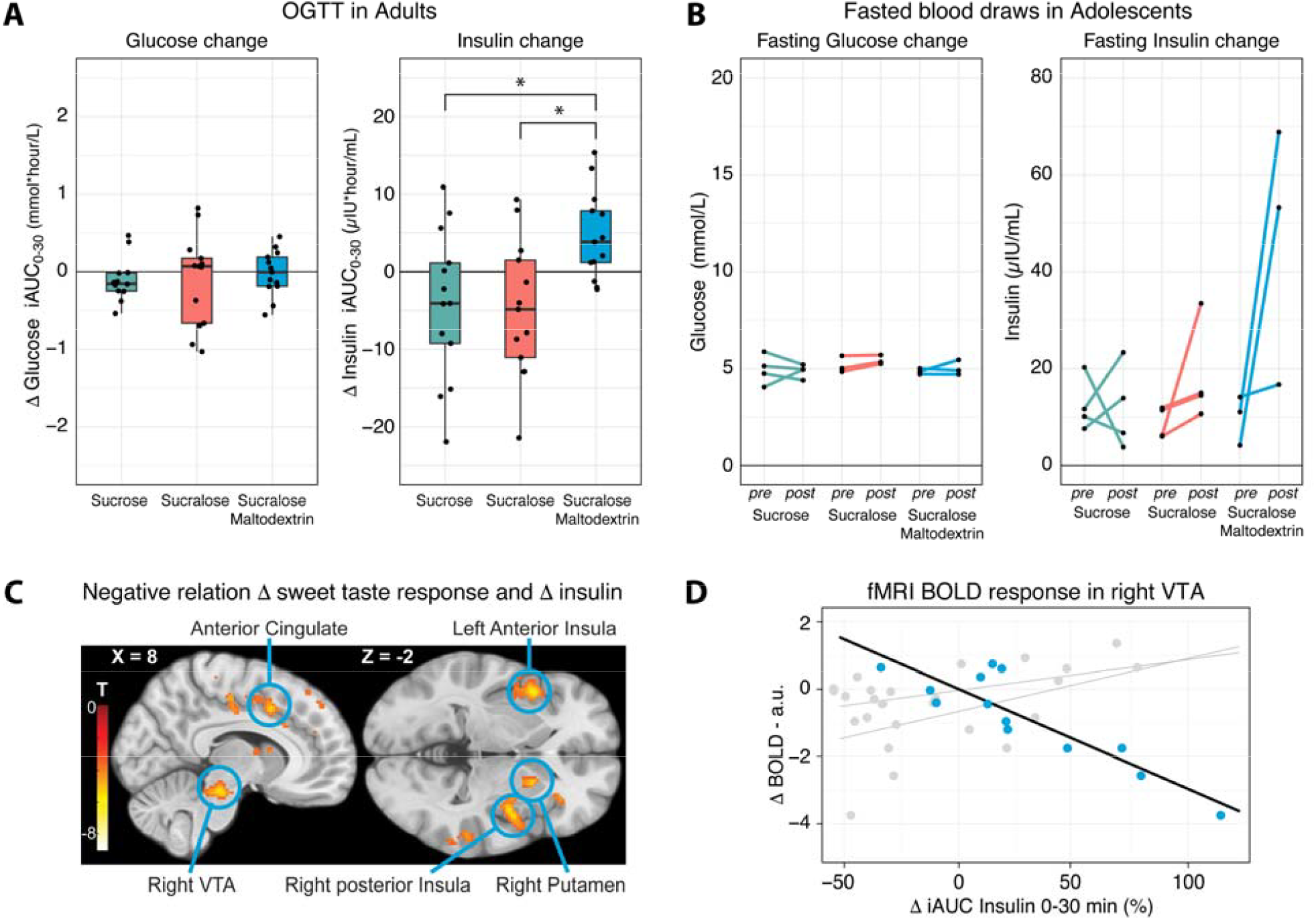
Changes in insulin sensitivity and brain response to sweet taste in humans. Changes in insulin sensitivity and brain response to sweet taste in humans (A) Relative change in first phase OGTT plasma glucose (left) and insulin (right) iAUC_0-30_ from pre to post beverage exposure. Post beverage exposure, insulin levels were significantly elevated in the sucralose plus maltodextrin group compared to the sucrose and sucralose groups (false discovery rate corrected *t* tests; both P=0.03). (B) Change in plasma glucose (left) and insulin (right) plotted per individual and per group in the adolescents study. The adolescent study was terminated because two participants in the sucralose plus maltodextrin group showed highly elevated insulin (and HOMA-IR) levels post beverage exposure. Permutation testing (n=1000) indicated that the difference scores of this group are significantly different from the sucrose and sucralose groups together (p=0.043). (C) Relative change in plasma insulin iAUC_0-30m_ from pre to post beverage exposure is significantly related to fMRI BOLD change in the sucralose plus maltodextrin young adults group during sucrose ingestion. The relation indicates that percent increase in blood insulin, decreases fMRI BOLD responses to sucrose in the anterior cingulate, left anterior insula, right substantia nigra/ventral tegmental area (VTA), right posterior insula and right putamen. All reported clusters are corrected for a cluster-wise FWE correction threshold of P<0.05. For visual purposes, brain maps are thresholded at p<0.001 (unc.). (D) Linear relationship between fMRI BOLD and relative change in plasma insulin iAUC_0-30m_ for peak voxels in the right VTA for the sucralose plus maltodextrin group (blue), and for the sucralose and sucrose groups (grey).

### Response to sweet, but not sour, salty, or savory taste in the ventral tegmental area, insula, putamen, and anterior cingulate cortex is inversely associated insulin sensitivity in the Combo group

To investigate the effect of beverage exposure on brain response to sweet taste and other basic tastes (sweet, sour, salty and umami – bitter was not used because of its lingering after-taste), we assessed blood-oxygen-level dependent (BOLD) changes in the brain using functional Magnetic Resonance Imaging (fMRI) in the adult study. We calculated fMRI-BOLD difference maps (post minus pre-beverage exposure) per taste on a single-subject level using mass univariate regression. At the group-level, we performed a mass univariate ANCOVA per basic taste to test whether brain response changed as a function of beverage exposure group while assessing the effect of insulin change as a covariate. Contrasting BOLD-difference maps between groups for each basic taste did not show any difference surviving a cluster-wise familywise error (FWE) correction threshold. However, regressing insulin iAUC difference scores on the BOLD-difference maps for sweet taste showed a strong negative relation in several limbic and mesolimbic areas (**Figure 2C**, **Table 1**) in the Combo group. In this group, the left anterior insula, right posterior insula, anterior cingulate, right ventral tegmental area (**Figure 2D**), right putamen, and several cortical areas in the superior temporal gyrus and postcentral gyrus showed a decreased fMRI-BOLD response to sweet taste as a function of iAUC. We found no association between insulin change and central processing of umami, salty, or sour taste nor any associations between insulin change and taste perception in the LCS and Carb groups.

**Table 1.** Negative relation between Δ insulin iAUC_0-30m_ and brain response to sucrose. The table shows the cluster-wise FWE corrected peak coordinates that show a decreased response when tasting sucrose as a function of increases in plasma insulin levels during the first 30 minutes of the OGTT in the sucralose plus maltodextrin adult group. The contrast was masked for grey matter only voxels. L: left; R: right; STG: superior temporal gyrus; SN: substantia nigra; VTA: ventral tegmental area; ACC: anterior cingulate cortex; SMA: supplementary motor area; PCG: postcentral gyrus.

| Region | Cluster |  | MNI {mm} |  |  |  |
| --- | --- | --- | --- | --- | --- | --- |
|  | p(FWE) | size k (2x2x2mm) | T | x | y | z |
| L Insula | < 0.001 | 385 | 8.95 | -46 | 12 | -8 |
| L Insula |  |  | 4.73 | -38 | 10 | -14 |
| L Rolandic operculum |  |  | 4.69 | -58 | 4 | 8 |
| R STG | < 0.001 | 901 | 6.1 | 50 | -20 | 6 |
| R STG |  |  | 6.02 | 66 | -26 | 6 |
| R pInsula |  |  | 5.54 | 38 | -4 | -2 |
| SN/VTA | < 0.001 | 265 | 5.54 | 8 | -26 | -18 |
| R Cerebellum |  |  | 5.49 | 16 | -36 | -20 |
| R Cerebellum |  |  | 5.29 | 22 | -42 | -26 |
| ACC | < 0.001 | 462 | 5.47 | -8 | 14 | 36 |
| ACC |  |  | 5.27 | 8 | 12 | 42 |
| SMA |  |  | 4.79 | 0 | -2 | 62 |
| R STG | 0.041 | 103 | 5.44 | 62 | -36 | 20 |
| L PCG | < 0.001 | 255 | 5.25 | -58 | -20 | 20 |
| L PCG |  |  | 4.69 | -60 | -20 | 34 |
| L PCG |  |  | 4.62 | -50 | -22 | 26 |
| L PCG | 0.026 | 115 | 4.68 | -30 | -44 | 58 |
| L PCG |  |  | 4.25 | -30 | -32 | 62 |
| L PCG |  |  | 3.67 | -24 | -28 | 72 |

### Taste intensity perception and preference is unaffected

To investigate the effects of LCS consumption on taste perception, we measured taste intensity ratings for sucrose, sucralose, citric acid (sour), sodium chloride (salty), monopotassium glutamate (umami), and sucralose+citric acid prior to each beverage exposure (7 times) across the two-week time period (**Figure 1**, **Figure S3 and Data S1**). We also assessed sweet concentration preference using a sucrose preference test pre and post beverage exposure. We found no differences in intensity perception or sucrose preference across the groups nor did we find an association between plasma insulin change and these measures (**Figure S3 and Data S1**).

### Summary Human Studies

Collectively, the findings from the two human studies refute the hypothesis that uncoupling sweet taste from caloric content causes metabolic dysfunction or decreases in the potency of sweet taste as a conditioned stimulus. Rather, the results reveal that metabolic dysfunction, coupled with reduced central sensitivity to sweet taste, occurs when an LCS is repeatedly consumed with, but not without a carbohydrate. Critically, while these findings fail to support the uncoupling hypothesis, they are nevertheless consistent with the results of the studies on which the hypothesis is based. More specifically, in these studies, LCS were added to yogurts that contained a number of nutrients including carbohydrates and thus metabolic dysfunction followed repeated simultaneous consumption of LCS and carbohydrates [14,18,20–22].

### Mouse Study

One possible mechanism by which an LCS could acutely alter carbohydrate metabolism is by influencing the glucose absorption rate [30,31]. Since altered glucose absorption would be expected to lead to changes in substrate utilization, we next turned to a mouse model where we could use calorimetric chambers to test whether consuming an LCS-sugar combination could alter nutrient partitioning (i.e. the substrate - lipids vs. carbohydrate - that the body uses for fuel). Young adult, fourteen-week-old C57BL/6J mice were individually housed in calorimetric chambers in a controlled environment with a 12:12 light-dark cycle with lights off at 19:00h. Mice were divided into 3 experimental groups and presented daily with a drink bottle containing a drinkable sweet solution at 14:00h for nine days. The three groups received sucrose (Sugar group: 0.25M, 0.337 kcal/ml), equi-sweet sucralose (LCS group: 0.425 mM, 0 Kcal) or equi-sweet sucrose plus sucralose (Combo group: 0.125 mM and 0.212 mM, respectively) containing half the caloric load of the sucrose solution (0.168 kcal/ml). We chose to mix sucralose with sucrose instead of maltodextrin because rodents show a strong taste preference for maltodextrins [32]. This also allowed us to determine if effects would be observed for carbohydrates other than maltodextrin. Volumes of 1.5ml were selected to be roughly equivalent to the human study where one 335ml beverage was consumed daily for nine days. Animals had *ad libitum* access to drinking water and food and every day, food intake (gram and kcal), liquid intake (ml), and locomotor activity (beam breaks/h) was measured every 15 minutes. To assess substrate utilization, we measured oxygen consumption (VO_2_) and carbon dioxide production (VCO_2_), and used these measures to determine Energy Expenditure (EE), Respiratory Exchange Rate (RER/RQ), and fat oxidation (FO).

### Nutrient partitioning is altered in mice when sucralose is consumed with, but not without sucrose

All groups fully consumed the solutions in a six-hour time window with no group differences in hourly consumption rate (**Figure 3A**); however, RER in mice consuming Sugar shifted more towards carbohydrate oxidation compared to mice in the LCS and Combo groups (false discovery rate corrected *t* tests; all P<0.02 between 14:00h and 17:00h and all P<0.024 between 15:00h and 18:00h, respectively) (**Figure 3B**). As this could be explained by either differences in nutrient metabolism or differences in nutrient intake, we next analyzed the RER adjusted for total caloric intake by calculating the hourly discrepancy between RER and the available macronutrients for oxidation based on chow and sucrose intake (food quotient, FQ) [33]. We found that, in the period with access to the sweet solutions, mice in the Combo group showed a larger discrepancy between FQ and RER (false discovery rate corrected *t* tests; all P<0.022 between 15:00h and 17:00h and all P<0.007 between 14:00h and 16:00h, for sucrose and sucralose groups, respectively) (**Figure 3C**). This indicates that there is a fundamental difference in the way food is metabolized in the context of consuming the solution with sucralose and sucrose. More specifically, substrate utilization (carbohydrate vs. lipids) differs between the groups despite similar absolute amounts of daily caloric intake. Consistently, the relationship between RER and caloric intake from the complete diet was different between the groups (group x total caloric intake interaction: F(2,138)=6.08, P=0.003) in the period with access to the sweet solutions (14:00-20:00, see **Figure 3A**) (**Figure 3D**). Post hoc tests revealed that mice in the Combo group differed from mice in both other groups (false discovery rate corrected *t* tests; β=0.053 VO2/CO2, P=0.037 and β=0.071 VO2/CO2, P=0.002, for the LCS and Sugar groups respectively) such that the shift from lipid to carbohydrate oxidation (i.e., substrate utilization) occurred at a higher energy intake in the combination group. This reflects an overall reduction in sensitivity to carbohydrates, which is consistent with our human experiments and with previous studies showing that glucose intolerance is associated with a metabolic shift towards fat oxidation [34]. Additional information on intake, activity, fat oxidation, and energy expenditure are reported in **Figure S4.**

**Figure 3.**
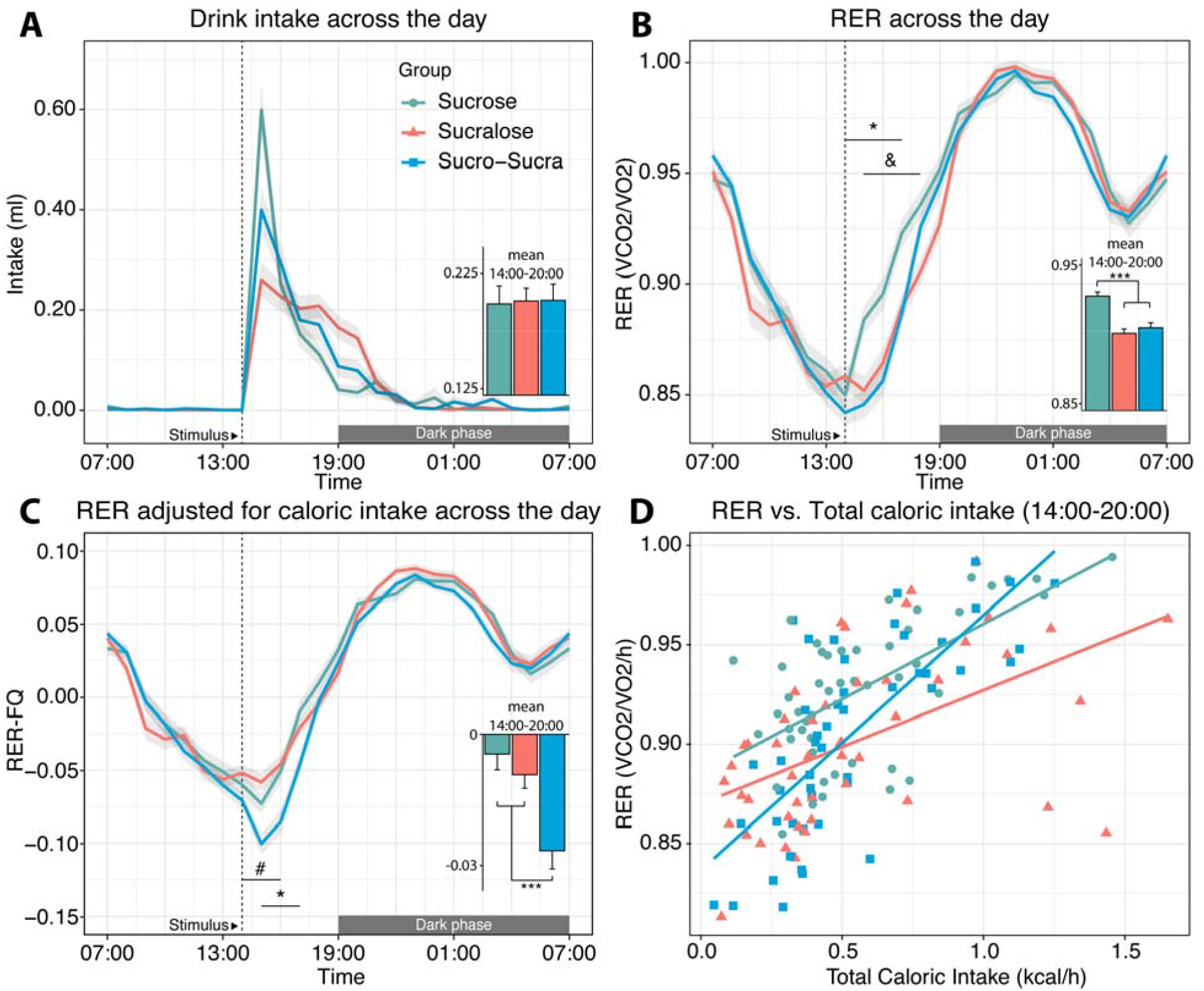
Metabolic changes in mice. (A) Hourly *ad libitum* drink intake of the presented 1.5ml solution across groups. We found no differences across groups in total intake, and average rate of intake between 14:00 and 20:00. (B) Hourly change in respiratory exchange ratio (RER) across groups. RER in the sucrose group deviated from the sucrose plus sucralose and sucralose groups (false discovery rate corrected across 3×24 *t* tests; all highlighted P<0.024). (C) Hourly change in the difference between food quotient (FQ) and RER. FQ-RER in the sucrose plus sucralose group deviated from the sucrose and sucralose groups (false discovery rate corrected across 3×24 *t* tests; all highlighted P<0.022). (D) Relation between RER and total caloric intake across groups during the period in which solutions were consumed (14:00h-20:00h, see A). The sucrose plus sucralose group deviated from the sucrose and sucralose groups (false discovery rate corrected across 3×24 *t* tests; P=0.037 and P=0.002, respectively). (A-C) Displayed values are mean values ± SEM error bands/bars. The onset of the solution presentation is highlighted with a dashed line at 14:00h. Bar graphs show the average difference across groups between 14:00h and 20:00h. (*) Sucrose plus sucralose vs sucrose, (#) sucrose plus sucralose vs sucralose, (&) sucrose vs sucralose. We found no group differences across all measures on a 24h time scale.

## Discussion

The results of our studies in humans and mice demonstrate that consuming sucralose with, but not without a carbohydrate rapidly impairs glucose metabolism. More specifically, in healthy human adults we observed reduced insulin sensitivity and blunted brain response to sucrose following consumption of seven 355ml beverages over two weeks, whereas no changes were observed following equal consumption of beverages with sucralose or sucrose alone. Likewise, in mice consuming 1.5 mls of a liquid containing sucrose and sucralose every day for seven days we observed reduced carbohydrate metabolism, whereas consuming equivalent volumes, but lower doses, of sucrose or sucralose alone had no effect. These results do not support the sweet uncoupling hypothesis. Rather, they suggest that sucralose consumption alters the metabolism of simultaneously consumed glucose to rapidly produce deleterious effects on metabolic health. Since the extent of this exposure is very likely experienced in a natural setting, our results provide evidence that LCS consumption contributes to the rise in the incidence of impaired glucose tolerance. They also indicate that the mechanism underlying this relationship involves acute LCS-induced alterations in glucose metabolism.

### The sweet uncoupling hypothesis

The current findings are consistent with the results of studies in rodents showing impaired glucose metabolism following repeated consumption of foods with added LCS (e.g., yogurt plus sucralose (Swithers refs); however, they refute the hypothesis that the impairment results from a decoupling of sweet taste with energy. First, in healthy adults who are non-regular consumers of LCS, repeated consumption of the sucralose beverage (i.e., group LCS), did not significantly influence glucose metabolism and produced no effects on brain or perceptual responses to sweet taste, despite being clearly rated as sweet-tasting and being decoupled from calories. Rather, in direct contradistinction, consuming a similarly sweet beverage containing the same dose of sucralose appropriately coupled to calories rapidly decreased insulin sensitivity. Second, the magnitude of the reduced insulin sensitivity was closely coupled to decreases in brain response to the sweet taste stimulus, whereas no main effects or correlations were observed with the responses to taste in the LCS and Malto groups. Although it is not possible to discern if this association results from altered central responses contributing to reduced insulin sensitivity or vice versa, it does suggest that central circuits, like peripheral glucose tolerance are altered by the exposure to the LCS only when it is coupled, rather than decoupled from calories. Third, the data from the adolescent study, though preliminary, are not only consistent with the adult findings, but also suggest that the negative impact of consuming sucralose and maltodextrin together is greater in youth, with 2 out of 3 participants in the combination group showing a clinically significant change in HOMA-IR. Finally, the rodent data align with the observations in humans, again pointing to an effect of LCS on metabolism only when consumed concomitant to a carbohydrate, in this case sucrose. This result is particularly impressive because the dose of sucralose was half that consumed by the LCS group and the caloric load from sucrose was half that consumed by the Sugar group. Nevertheless, it was the Combo group that displayed reduced carbohydrate oxidation in the context of similar overall liquid and food intake. Collectively, the results from all three experiments are consistent and lead to the conclusion that consumption of the LCS sucralose with, but not without a carbohydrate, produces metabolic dysfunction.

### Possible mechanisms

Our findings argue that uncoupling sweet taste from calories cannot be responsible for associations that are observed between LCS consumption and impaired glucose metabolism. Rather they point towards a mechanism that operates when LCS and carbohydrate are consumed concurrently. LCSs, including sucralose, bind to T1R2/T1R3 sweet taste receptors that are expressed in a variety of tissues including the oral cavity, intestine, liver, pancreas and brain [35]. Activation of sweet taste receptors expressed in the intestine by LCSs produce up-regulation of sodium/glucose co-transporter SGLT-1[31], which plays a role in glucose absorption and are implicated in the ability of dietary supplementation of LCS in piglets to increase weight gain [36]. The binding of LCS to intestinal taste receptor cells may also influence absorption via the translocation of GLUT2 [15,30,37]. Considering the current study, maltodextrin is quickly metabolized into glucose, which would then be available to bind to intestinal taste receptor cells. Simultaneous binding of maltodextrin-derived glucose and sucralose could therefore increase glucose transport (by SLT-1 and/or GLUT2) beyond optimal levels for the amount of glucose present, resulting in acutely perturbed glucose homeostasis. Consistent with this possibility, in obese, but glucose tolerant humans, consuming sucralose compared to water prior to an OGTT, results in higher peak plasma glucose concentrations, increased insulin concentration and AUC, and decreased insulin sensitivity (Pepino et al., 2013). Importantly, this work excluded individuals who self-reported consuming more than the equivalent of 1 diet soda per week. In contrast, studies that have not excluded regular users have failed to find effects of LCS consumption on glucose metabolism [38–41]. However, as suggested by Pepino and colleagues, negative results would be expected if regular consumption of LCS impaired glucose tolerance. Our findings align with this proposal. Like Pepino and colleagues, we excluded individuals who self-reported consuming LCS more than three times per month. Further, examination of the TLFB questionnaire data indicated that participants consumed an average of 260 mls of diet drinks per week - which is less than three 355ml bottles per month. In this case, our intervention (seven, 355, ml bottles in two weeks) clearly increased consumption above baseline levels and as would be predicted from the acute effects observed by Pepino and colleagues, resulted in a longer term decrease in insulin sensitivity. Critically, in our study sucralose was not consumed prior to the OGTT. Therefore, the observed decrease in insulin sensitivity must be attributed to a chronic effect of consuming the Combo beverage on glucose tolerance.

It is also possible that altered substrate utilization plays a role in the development of glucose intolerance. We observed that mice consuming the sucralose plus sucrose solution had altered substrate utilization favoring fatty acid over carbohydrate oxidation. More specifically, shifting from lipid to carbohydrate oxidation (i.e., substrate utilization) required higher energy intake in the sucrose plus sucralose group compared to the other groups. As in our studies in humans, this shift reflects a decreased sensitivity to carbohydrates, which could over time promote increased adiposity and decreased glucose tolerance. Interestingly, similarly altered substrate utilization is well known to occur in T2D, but it is unknown whether this precedes or follows insulin resistance [42]. Our results suggest that diet can rapidly alter substrate utilization and may therefore precede impaired metabolism. In future work, it will be important to determine whether consuming an LCS with a carbohydrate produces similar effects in substrate utilization.

An alternative, or possibly adjunct possibility, is that change in insulin sensitivity is regulated by diet-induced changes in dopamine signaling. Manipulating central dopamine circuits can influence peripheral insulin sensitivity [43]. In the adult study, peripheral changes in insulin sensitivity were strongly associated with central changes in BOLD responses to sweet taste in the insular taste cortex as well as dopaminergic source and target areas. This effect did not result from a general influence on taste processing since associations were not observed with salty, sour or umami tastes. It is also unlikely to reflect sensory processing of taste since perception was unaltered, or satiety signaling because the amount of each substance consumed during the course of the fMRI study was negligible (e.g. less than 20 mls per stimulus). It is also unlikely to reflect a learned decrease in the nutritive value of sweet taste since no associations were observed with the LCS alone condition. Rather, we suggest that consuming the Combo beverage decreased either peripheral or central insulin sensitivity, which then led to blunted dopamine response to carbohydrates reflected in decreased BOLD response to sweet taste. This possibility is consistent with a number of related findings. In humans, peripheral insulin resistance is associated with decreased insulin induced activation in the ventral striatum [44] and correlates with dopamine type 2 receptor availability [45]. In fruit flies, chronic exposure to sucralose alters the equivalent of insulin and dopamine systems leading to hyperactivity, insomnia, glucose intolerance, enhanced sweet taste perception and increased food intake [46]. In rodents, diet-induced reduced central insulin sensitivity is known to blunt Akt (protein kinase B) induced mobilization of the dopamine transporter (DAT) leading to blunted striatal response to amphetamine evoked dopamine response [47]. We therefore speculate that blunted brain response is a consequence rather than a cause of insulin resistance. However, the alternative cannot be ruled out.

### Resolving the inconsistencies in the literature

As mentioned above, although our results fail to support the uncoupling hypothesis, they are nevertheless consistent with the results of the studies on which this hypothesis is based since LCS was added to carbohydrate-containing foods and therefore parallel our Combo groups. In many rodent studies reporting a negative impact of LCS on metabolism, LCS (e.g., saccharin, aspartame, sucralose, AceK) were either added to a carbohydrate or a carbohydrate-containing yogurt stimuli ranging from 0.4 to 0.6 kcal/g [18,21,22,48,49]. Similarly, in human randomized control trials (RCTs) reporting that LCS consumption impairs metabolism, LCS were consumed concomitant to carbohydrates [50–53]. Critically, when study protocols promote consumption of LCS alone or in capsules at home during meal times, studies fail to find a negative impact on metabolism [54–57]. This suggests that LCS may have different effects depending on how they are consumed, with greater likelihood for impairment when LCS are provided in conjunction with carbohydrate.

Another important factor, as proposed by Pepino and colleagues is that results may depend on individual factors like prior experience consuming LCS. More specifically, including individuals who are regular users of LCS may bias towards negative findings because these individuals might already be affected and would therefore be less likely to show a change upon additional limited small exposures. A further important issue is that LCS are biochemically heterogeneous and have diverging bioactive effects. Sucralose is the most commonly used LCS, but there are several other in frequent use and possessing different pharmacokinetics (i.e. absorption, distribution, metabolism and excretion) [58]. Several studies (reviewed in [59]) suggest that the effects of LCS on glucose transporters and subsequent absorption are strongest for Ace-K and weak, or absent for aspartame. For example, sucralose and Ace-K, but not aspartame, increase SGLT-1 mRNA expression, which correlates with absorption rate [31]. In addition, Ace-K and sucralose, but not aspartame, increase insulin secretion [60,61]. One reason why aspartame may produce less effects on incretins and glucose absorption is that it is rapidly metabolized in the small intestine and would therefore have less opportunity to bind to taste receptor cells or glucose transporters. Given the potential for insights into mechanisms as well as importance for health, future work should focus on comparing different categories of LCS within the same study.

### Summary and Implications

The results from our studies demonstrate that LCS consumption produces metabolic dysfunction when it is consumed with, rather than uncoupled from, a carbohydrate. This implies that (a) carbohydrate metabolism is altered in the presence of the LCS sucralose and (b) that this alteration leads to decreases in peripheral and central sensitivity to sugar and sweet taste. Of particular relevance to the potential significance of this work, the metabolic changes observed following a very limited exposure that almost certainly occurs in freely living humans - especially if one considers the consumption of a diet drink along with a meal. This raises the possibility that the combination effect may be a major contributor to the rise in the incidence of type two diabetes and obesity.

## Acknowledgements

We would like to acknowledge those volunteers who participated in our study, the MR technicians, as well as several colleagues, including Barry Green, Ivan de Araujo and Wolfgang Meyerhof for insightful discussions and comments on design, interpretation or prior drafts of this manuscript. Funding: NIH NIDCDR01 DC006706-06A1.

## Author Contributions

Conceptualization: DMS, SL, MGV, BPP

Data Curation: JRD, BPP, RD

Formal Analysis: JRD, RD, BPP, MGV

Funding Acquisition: DMS

Investigation: BPP, RD, YN, PCV

Methodology: DMS, JRD, RD, MGV, SL

Project Administration: JRD, BPP, RD, MGV, SL, DMS

Resources: SL, DMS

Software: JRD, MGV, RD

Supervision: SL, DMS

Validation: JRD, BPP, RD

Visualization: JRD, RD

Writing – Original Draft Preparation: JRD, DMS

Writing – Review & Editing: JRD, BPP, RD, MGV, YN, PCV, SL, DMS

Competing interests: Authors declare no competing interests.

## STARS METHODS

### CONTACT FOR REAGENT AND RESOURCES SHARING

“Further information and requests for resources and reagents should be directed to and will be fulfilled by the Lead Contact, Dana Small.

### EXPERIMENTAL MODEL AND SUBJECT DETAILS

#### Experiments in Human Subjects

Participants were recruited through advertisements around Yale University and the greater New Haven area. Participants were screened either over the phone or through an online screening form. Participants aged 23-45 years were assigned to groups matched for sex, age, and BMI. Exclusion criteria were obesity (BMI>30), frequent NNS-user (self-report of > 3 times a month), history of psychiatric disorders, eating disorders or head injury with loss of consciousness, being on a diet, alcoholism, tobacco or drug use, use of daily medication other than monophasic birth control, chemosensory impairments, lactose intolerance, food allergies, and ineligibility for an fMRI scan. The study was approved by the Yale Human Investigations Committee and all participants provided written informed consent at the start of their first lab visit.

#### Experiment 1

55 adult participants were recruited, 6 dropped out during the experiment, blood sampling during at least one full OGTT failed for 4 participants, and 6 participants later revealed in a timeline follow back measurement that they were regular users of NNS. Perceptual, blood, and brain data of these subjects were discarded before data analysis. Data analysis was performed on data from 39 adult subjects (13 per group; 21 women; mean age 27.79 ± 3.96; mean BMI 23.72 ± 3.13).

#### Experiment 2

Participants, aged 13-17 years were recruited similarly to the adult study. The study was approved by the Yale Human Investigations Committee and all participants provided written informed assent at the start of their first lab visit together with their parents who provided parental consent. Before the Yale Human Investigations Committee advised study termination, 17 adolescents were recruited, 2 refused blood sampling, 3 dropped out, and 1 group assignment was lost in a software crash. Data analysis was performed on data from 11 adolescents (8 female; mean age 15.95 ± 1.37; mean BMI 22.13 ± 3.63).

#### Experiment 3

Three groups of 14-week-old mice (n=8) C57Bl/6J (from Janvier, Le Genest St Isle, France) housed in stainless steel cages in a room maintained at 22 ± 1°C with light from 7am to 7pm. Food (3.236 kcal/g. Safe, Augy, France) and water were given *ad libitum* unless otherwise stated. Animals were caged individually one week prior to experiments. All animal experiments were performed with approval of the Animal Care Committee of the University Paris Diderot-Paris 7 and according to European directives.

## METHODS DETAILS

### Experiment 1 & 2

#### General Procedure

After screening and acquiring informed consent, participants were assigned to either the sucrose, sucralose, or sucralose plus maltodextrin group. Participants completed a nutrition questionnaire (NQ) to screen for inclusion and exclusion criteria, two sucrose preference tests (pre and post), a training session, anthropometric measurement, two Monell forced-choice sweet taste preference tests (M-STP; pre and post), two timeline follow back sessions (TLFB; pre and post), two blood sampling sessions (adults completed OGTTs) while adolescents completed fasting blood draws), two fMRI scanning sessions (pre and post), seven psychophysiological measurements, seven beverage exposures, and a debriefing (**Figure 1**). Exposure sessions were conducted on separate days within 2 weeks.

On the first pre-exposure session, the exposure sessions, and last post-exposure session, participants arrived at the lab after a 1h fast. For blood sampling, participants arrived after a 10-12 hour overnight fast. For fMRI scans, participants were instructed to arrive neither hungry nor full on each scan day.

The TLFB Questionnaire, sweet taste preference tests, and Psychophysiological measurements are reported in **Data S1**.

#### Training session & anthropometric measurement

A pregnancy/toxicology screening was performed, and height was measured using a stadiometer. Body weight and body fat percentage were measured using the BodPod body composition tracking system [62] in minimal attire (spandex shorts and sports bra for women). Following anthropometric measures, participants were trained to make computerized ratings of their internal state as well as the perceptual qualities of various stimuli on computerized scales. Internal state ratings were made up of a series of adapted cross-modal gLMS consisting of a 100mm vertical line scale with the labels “barely detectable” at the lower endpoint and “strongest imaginable sensation” at the upper endpoint [63,64]. Participants were instructed to rate the intensity of their feelings of hunger, fullness, thirst, anxiety, and need to urinate. The perceptual qualities of real and imagined stimuli consisted of ratings of their overall intensity, liking, and wanting to eat. Liking was measured using a labeled hedonic scale consisting of a 100 mm vertical line scale with the labels “most disliked sensation imaginable” at the lower anchor point, “most liked sensation imaginable” at the upper anchor point, and “neutral” in the middle. Wanting to eat was rated on 200 mm visual analog scales labeled on the left with “I would never want to consume this” and “I would want to consume this more than anything” on the right. Participants also rated the perceptual qualities of basic tastes (sucrose, 0.56M; citric acid, 18mM; NaCl, 0.32M; quinine, 0.18mM, and MPG (100mM) alone and when combined as binary taste mixtures (sucrose-citric acid, sucrose-quinine, sucrose-MPG, citric acid-NaCl and NaCl-quinine). Participants rated the sweetness, sourness, saltiness, bitterness, and umami intensity of each taste using the gLMS. In addition, the experimenter assisted the participant in completing a TLFB questionnaire in which all beverages (besides water) consumed over the previous 14 days were written down, including brands and amounts. This questionnaire ensured that participants were not regular users of NNS.

Lastly, participants underwent an fMRI training simulation to familiarize themselves with the paradigm, learn to remain still in the scanner, and reduce anxiety on the day of the scan (see fMRI sessions for more details).

#### Beverage exposure sessions

##### Stimuli

Exposure beverages contained 355ml of a novel-flavored equi-sweet solution. Beverages contained either 0.06g sucralose (0 Kcal, Sigma-Aldrich Inc. MO, USA), 30.38g sucrose (120 Kcal), or 0.06g sucralose and 31.83g of the non-sweet carbohydrate maltodextrin (120Kcal) (Maltodextrin, FCC, M1083, Spectrum Chemical Mfg. Corp.). Beverages were colored and flavored according to the preference of each participant. Participants could choose any color (1-3 drops; McCormick & Co, Inc. MD, USA, Assorted food color & egg dye: Red, Yellow, Green, Blue; McCormick & Co, Inc. MD, USA, NEON! Food color & egg dye: Purple, Green, Pink, Blue) and between an Aloe Vera or Papaya flavor (0,355ml; Aloe Vera, Bell Labs, ID#:141.31480; Papaya, Bell Labs. ID#102.82506).

##### Procedure

Subjects were invited seven times to the lab across a time span of two weeks. Subjects were first asked to perform a psychophysiological measuring perceptual taste thresholds (**Data S1**). Subsequently subjects received their respective exposure beverage and were asked to finish the drink within five minutes.

#### Blood sampling sessions

In the human young adult study, we performed OGTTs. Upon arrival, an indwelling intravenous line was placed by an experienced nurse or phlebotomist, followed by a 20 min rest period in order to limit any stress of the catheter placement on the blood measures. Participants were asked to fully consume (within ∼2min) an orange-flavored drink containing 75 g of dextrose (10 oz, Trutol, VWR, Radnor, PA). Blood was drawn at 0, and then 15, 30, 60, 90 and 120 min post-drink and immediately placed into tubes. For adolescents, only one fasting blood sample for measurement of HOMA-IR was taken pre and post exposure.

#### fMRI scans

##### Stimuli and Delivery

Taste stimuli for fMRI scans included a sweet sucrose solution (0.32 M), a sour citric acid solution (0.0056 M), a salty sodium chloride solution (0.14 M), an umami monopotassium glutamate solution (68 mM), and a tasteless and odorless solution.

A custom-designed gustometer was used to deliver liquid stimuli. This system has been successfully used in past fMRI studies [65–67]. This gustometer system is a fully portable device that consists of a laptop computer that controls (via a 9-pin serial adaptor and telephone wiring) up to 11 independently programmable BS-8000 syringe pumps (Braintree Scientific, Braintree, MA) to deliver precise amounts of liquids to subjects lying in the mock or real scanner at precisely timed intervals and durations. The pumps, which infuse liquids at rates of 6-15 mL/min, are controlled by programs written using Matlab 7.11 (MathWorks Inc., Sherborn, MA) and Cogent2000 v1.25 (Wellcome Department of Cognitive neurology, London, UK). Each pump holds a 60 mL syringe connected to a 25-foot length of Tygon beverage tubing (Saint-Gobain Performance Plastics, Akron, OH) with an inside diameter of 3/32”. All tubing terminates into a specially designed Teflon, fMRI-compatible *gustatory manifold* (constructed in the Pierce Laboratory Electronics and Machine Shop), which is anchored to the MRI headcoil and interfaces with the subject. This set-up is depicted below in a close-up (**Figure S2A**), with the subject in the mock scanner (**Figure S2 and Data S1**).

The gustometer mouthpiece or “manifold” was designed to deliver up to 11 taste solutions and one tasteless rinse. All tastants and rinses pass through 1-mm channels that converge at a central point at the bottom of the manifold for delivery to the tongue tip. To prevent the subject’s tongue from coming in contact with the 1mm holes, and to ensure the liquids flow directly onto the tongue, a short silicone tube is attached to the outflow point under the 1-mm holes. The subject holds the silicone tube between their lips and teeth, and the tip of the tongue rests up against the lowest point of the tube. A large vent hole prevents subjects from drawing or sucking the stimulant through the manifold at uncontrolled times or rates. Tactile stimulation is held constant across all events (i.e. delivery of the different tastants and the tasteless solutions) by the use of converging outflow – so that the liquid arrives at the same location for each stimulus. The gustometer manifold is mounted by rigid tubing onto an anchoring block that clamps onto the front of the head coil. The anchor height and horizontal positions are adjustable via two knobs accessible to the subject and the experimenter to achieve the most comfortable position. The manifold is then locked in place for the duration of the scanning run. This setup has previously been described by Veldhuizen et al [65].

All scans were scheduled between 10am and 3pm. Sweet, sour, salty, umami, and tasteless stimuli were presented in a block design across two functional imaging runs. During each block, 4 to 8 uncued taste stimulus presentations were presented with a volume of 0.75ml delivered over 2s followed by a 7s swallowing period. Each taste block was presented four times and block length varied between 36 to 54 seconds. The order of blocks was counterbalanced across subjects. Each taste block was followed by a rinsing period (0.75 ml deionized water over 2 seconds). Blocks were separated with a 10 second rest-period.

MRI scans were performed using a Siemens 3.0 Tesla TIM Trio scanner at Yale University Magnetic Resonance Research Center equipped with a 32-channel head coil. A T1-weighted 3D MPRAGE whole brain image was acquired for anatomical reference. Acquisition parameters: TR/TE: 1900ms/2.52ms; flip angle: 9°; FOV: 250; matrix: 256 × 256; slice thickness: 1 mm; number of slices: 176, scan duration = 4:18 min. T2*-weighted functional brain images were acquired using a multiband susceptibility-weighted single-shot echo planar imaging sequence. Acquisition parameters: TR/TE: 1000ms/30ms; flip angle = 60°; FOV = 220 mm; matrix = 110 × 110; slice thickness=2 mm; and acquisition of 60 contiguous slices. Slices were acquired in an interleaved order. Each functional taste run lasted for 12:02 minutes. The first 2 volumes of each run allowed the MR signal to equilibrate (“dummy images”).

#### Post-test sessions

At the end of the experiment, participants were invited once more to perform the second M-STP and to fill in another TLFB questionnaire to measure whether participants changed their NNS consumption. Subsequently, participants were debriefed about the goal of the study.

### Experiment 3

#### General Procedure

Mice were individually housed 7 days before entering the calorimetric chambers, accustomed to their new cages and drinking bottle. Every day mice were also presented with a 1.5 mL drinkable solution of either sucrose (0,25M, 0.337 kcal/mL), sucralose (0.425 mM) or a mix of sucrose and sucralose (0.125 mM and 0.212 mM respectively, 0.168 kcal/mL) at 2 pm, the solution bottle was refilled the following day. Once accustomed to the drinking solution with no sign of phobia, animals were installed in calorimetric chambers. All animals were acclimated to the chambers containing a calorimetric device (Labmaster, TSE Systems GmbH, Germany) for 48 h before experimental measurements.

#### Indirect calorimetry

All animals were acclimated to the chambers containing a calorimetric device (Labmaster, TSE Systems GmbH, Germany) for 48 h before experimental measurements. We recorded food intake (grams), experimental drink intake (ml), activity (beambreaks/h), oxygen consumption (VO2), and carbon dioxide (VCO2) production in 15-minute time intervals. Whole lean tissue mass was extracted from the EchoMRI (Whole Body Composition Analysers, EchoMRI, Houston, USA) analysis as previously described Joly-Amado *et al.* [68]. From the available data, we calculated energy intake in kcal, respiratory exchange ratio (RER, VCO2/VO2), energy expenditure (EE), fatty acid oxidation (EE x (1-RER/0,3) [69], and the oxidation ratio (see below).

## QUANTIFICATION AND STATISTICAL ANALYSIS

### Experiment 1 & 2

#### Blood sampling sessions

Blood samples were centrifuged, frozen immediately and stored at –80°C until analysis. Plasma glucose was analyzed using the YSI Life Sciences 2300 STAT PLUS Glucose and L-Lactate Analyzer. Plasma insulin (sensitivity: 0.1817 mU/L (1.09 pmol/L)) was measured using the Eagle Bioscience insulin ELISA kit (Eagle Bioscience, Nashua, NH). All samples were analyzed in duplicate. The sample average was used for statistical analysis.

Statistical analysis was performed in R. For the adult trial, 8 out of 480 blood samples were missing at random. Missing values were imputed separately for plasma glucose and insulin using a Principal Components Analysis model [70] available in the package missMDA (version 1.13). We used this form of imputation as it uses the interrelation across measurement time points for insulin and glucose curves, respectively. Subsequently, incremental area under the curve (iAUC) values for plasma insulin and glucose were calculated for the first 30 minutes of the OGTT using the auc-function in the MESS-package (version 0.5.2). We then calculated absolute and relative iAUC difference scores (%). To test for group differences in insulin levels, ΔInsulin iAUC_0-30m_ was entered in a linear model as dependent variable while group ID was entered as independent variable to test for group differences. In a similar model, we tested for differences in glucose levels by entering ΔGlucose iAUC_0-30m_ as a dependent variable while group ID constituted the independent variable. For completeness, we also investigated the results for the complete OGTT AUC using an identical statistical analysis procedure. Additionally, we calculated the Matsuda index, a measure of whole-body insulin sensitivity [71] and Hepatic insulin resistance index [72].

Matsuda index:

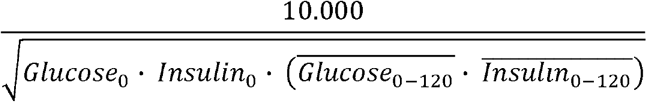

Hepatic insulin resistance index:

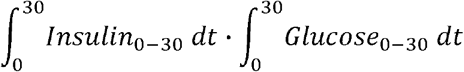

We found no group differences for the Matsuda index. However, there were differences among the groups for the Hepatic insulin resistance index (F(2,36)=3.79, p= 0.03). The effects are similar to the iAUC_0-30_ results. Furthermore, we found a difference between the sucralose and sucralose plus maltodextrin group for the OGTT AUC_0-120_ and incremental AUC_0-120_. The results are given in **Figure S1** and **Table S2**.

#### fMRI scans

fMRI data were analysed using SPM12 (v6906, Wellcome Trust Centre for Neuroimaging, http://www.fil.ion.ucl.ac.uk/spm) running in Matlab 2016b (The MathWorks Inc., Natick, MA). The first 2 dummy images from each functional run were removed. Subsequently, functional images from both visits were realigned, co-registered to the T1-weighted anatomical image acquired during the first visit, normalized to MNI space, and smoothed with a 6 mm FWHM kernel. For first-level statistical analysis, we constructed mass-univariate general linear regression models for each participant. The regressors included: 1) conditions ‘Sweet’, ‘Sour’, ‘Salty’, ‘Umami’, and ‘Tasteless’, and 2) the realignment parameters and their first derivatives as covariates[73]. The task-related regressors were convolved with the canonical hemodynamic response function (HRF) and a high-pass filter of 128 seconds was applied.

Prior to the group analyses, we calculated difference maps [post scan - pre scan] per taste condition. On group-level we performed a mass univariate ANCOVA on these difference maps per basic taste to investigate whether brain responses changed as a function of beverage exposure group. To test whether changes were attributed to dysregulation in insulin signaling, we entered ΔInsulin iAUC_0-30m_ as covariate. FWE correction for the mass univariate analyses was performed on a cluster level.

### Experiment 3

#### Calorimetry

We recorded food intake (grams), experimental drink intake (ml), activity (beambreaks/h), oxygen consumption (VO2), and carbon dioxide (VCO2) production in 15-minute time intervals. Whole lean tissue mass was extracted from the EchoMRI (Whole Body Composition Analysers, EchoMRI, Houston, USA) analysis as previously described Joly-Amado *et al.* [68]. From the available data, we calculated energy intake in kcal, respiratory exchange ratio (RER, VCO2/VO2), energy expenditure (EE), fatty acid oxidation (EE x (1-RER/0,3) [69], and the oxidation ratio (see below).

Analysis of the data was performed in R 3.5.1 (2018-07-02). First, we changed the temporal resolution from 15-minute time bins to 1h time bins using the eXtensible Time Series (xts) package (v0.11-1).

In a state of energy balance, macronutrient oxidation should match macronutrient intake. In other words, the proportion of macronutrients oxidized by the body reflected in the respiratory exchange ratio (RER) should equal the proportion of dietary macronutrients available for oxidation, reflected in the food quotient (FQ). Many studies have shown that when people are in energy balance, 24-h mean RER equals 24-h FQ [74–76]. The difference between the RER and the FQ expresses the oxidation ratio and reflects a discrepancy between the macronutrients available for oxidation and the effective body oxidation (use of substrate). Following Jéquier *et al* (1987) [33], FQ can be extrapolated from the respiratory quotients of protein (0.80), fat (0.71) and carbohydrate (1.00). As there was no change in chow diet and sucralose contains no carbohydrates, FQ was constant in the sucralose group. For the sucrose and sucralose plus sucrose groups we updated FQ based on hourly food and drink intake.

Statistics were performed using LMM (LMMs, package LME4, version 1.1-18-1) [77]. P-values were calculated using the Satterthwaite’s approximation for degrees of freedom, provided in the package lmerTest (version 3.0–1) [78]. Differences across groups were statistically tested per 1-hour time bin across a 24h time span. For each dependent variable, three tests were performed in each time bin; 1) sucrose vs sucralose, 2) sucrose vs sucralose+sucrose, and 3) sucralose vs sucralose+sucrose, resulting in 24 x 3 = 72 comparisons. All reported p-values are adjusted using the false discovery rate across these 72 comparisons.

#### Data and software availability

Raw MRI data:https://openneuro.org/ pending

Statistical maps of the human brain: https://neurovault.org/ pending

All other data: https://data.mendeley.com/ pending

