## Supplementary Information for "Short-term consumption of sucralose with, but not without, carbohydrate impairs neural and metabolic sensitivity to sugar"

**Data S1. TLFB Questionnaire, Sweet taste preference** **and Psychophysiological measurements**

**TLFB Questionnaire**

*Procedure*

The TLFB (*5*) was used to estimate LCS beverage intake as a proxy for LCS intake. The TLFB was administered by the experimenter who asked participants to retrospectively report their daily beverage intake for the 14 days prior to the interview. The TLFB interviewer used an annotated calendar, started with the day of the interview and asked the participant to report all consumed beverages of that day and estimate the consumed volume by using plastic replicas of different types of drinking glasses as examples. This procedure was repeated 14 times in reverse chronological order. The TLFB was performed pre and post beverage exposure.

*Analysis and results*

In addition to measuring LCS consumption using the TLFB to control exclusion criterion, we tested whether our experimental manipulation affected sugar sweetened beverage (SSB) consumption outside of the experiment. To test for (a) any group differences in SSB consumption, (b) group x time (pre vs post exposure) differences in SSB consumption, and (c) any associations between ΔInsulin iAUC_0-30m_ and SSB consumption, we performed two linear mixed models (LMMs) using package LME4 (version 1.1-7) (*6*). Subsequent statistical tests on the LMMs were performed using the Satterthwaite’s approximation for the degrees of freedom, provided in the package lmerTest (version 2.0-11, <http://cran.r-project.org/package=lmerTest>) (*7*). As dependent variables, we entered drinking days and amount consumed for the first and second model, respectively. As independent variables, we entered a group x time interaction and a group x ΔInsulin iAUC_0-30m_ interaction. Reported model results are based on the type III Analysis of Variance Table of the LMMs.

As drinking days and consumed volume (in ml) are count variables, we took extra care in optimally fitting the LMMs to the data. As LMMs assume that the residual error distribution of the model is normally distributed, we excluded outliers with a residual error greater than 2.5 standard deviations from 0 using the ‘romr.fnc’-function, available in LMERConvenienceFunctions (version 2.5) (*8*, *9*). Subsequently, we refitted the models to the trimmed data if necessary. Residual errors in the LMM fitted on drinking days contained no outliers, whereas 4 outliers (5.1%) were removed based on the LMM fitted on consumed volume. The resulting residual error distributions of both models did not deviate from normality (Shapiro-Wilk W=0.982, p=0.34 and W = 0.9924, p-value = 0.94, for the drinking days and consumed volume model, respectively).

Although the mixed-effects repeated-measures ANOVA table indicated a difference in the total number of SSB drinking days (F(2,32)=4.74, p<0.05), the total amount consumed did not differ across groups (F(2,31)=2.17, p=0.13), nor did we find any group x time interactions (F(1,36)=0.11, p=0.74; F(1,34.64)=0.30, p=0.58 for drinking days and total amount, respectively). Furthermore, we found a significant group x ΔInsulin iAUC_0-30m_ interaction for SSB drinking days (F(2,32)=3.94, p<0.05). Post hoc contrasts indicated a significant negative association between SSB drinking days and ΔInsulin iAUC_0-30m_ in the sucrose plus maltodextrin group (β= -0.0095, t(32)=-2.45, p<0.05). This association was significantly different from the sucralose group (β= 0.012, t(32)=-2.78, p<0.01).

Together these results indicate that we found no evidence that our experimental manipulation affected SSB consumption outside the experiment. In contrast, we did find that participants who drink SSBs on a more regular basis had a lower change in plasma insulin between pre and post OGTT measurement in the sucralose plus maltodextrin group.

**Sweet taste preference**

*Stimuli*

We used the sucrose concentrations 3% - 0.09M, 6% - 0.18M, 12% - 0.35M, 24% - 0.70M, and 36% - 1.05M.

*Procedure*

The M-STP (*6*) task was performed pre and post beverage exposure to investigate psychophysiological changes in sweet taste preference. Subjects sampled (sip-and-spit) two cups containing 10 ml sucrose dilutions and chose the one they preferred in a forced choice setting. If the subject selected the higher concentration, the next two cups presented was the selected beverage plus the next highest concentration. If the subject selected the weaker concentration, the next two presented cups were the weak solution plus the next lowest concentration. Forced choices were presented until the subject selected the same concentration two times in a row. The procedure was completed twice; starting at 3% and 36% sucrose, respectively.

*Analysis and results*

Statistical analysis was performed in R (version 3.1.3, 2015-03-09). First, we made the step size in the concentration range equal by converting the preferred concentrations to a log scale (correlation between log concentrations and linear scale, r = 0.996). The preferred level of sucrose was estimated by averaging the two chosen concentrations (on log scale) for the pre and post measurement, separately. Subsequently, difference scores were calculated (post minus pre beverage exposure) and submitted to a type III Analysis of Variance (ANOVA) model to investigate changes in sweet taste preferences across groups and to investigate whether changes in sweet taste preference were associated with changes in insulin sensitivity. ΔLog sucrose preference was entered as dependent variable whereas group * ΔInsulin iAUC_0-30m_were entered as independent variables. Results are reported in the Supporting information.

We found no differences across groups (F(2,32)=0.16, p=0.85). We found no association between changes in sucrose preference and changes in insulin sensitivity (F(1,32)=0.05, p=0.82), nor did we find a group * ΔInsulin iAUC_0-30m_ interaction (F(2,32)=2.08, p=0.14). These results indicate that we found no evidence that our experimental manipulation affected sucrose preference. Though, we are unable to rule out that the M-STP was not sensitive enough.

**Psychophysiological measurements**
*Stimuli*

Taste stimuli included a sweet sucrose solution (0.32 M), a sour citric acid solution (0.0056 M), a salty sodium chloride solution (0.14 M), an umami monopotassium glutamate solution (68 mM), a sweet sucralose solution (0.588 mM) and a sweet and sour sucralose + citric acid solution (0.588 mM + 0.009 M).

*Procedure*

Prior to each beverage exposure, we measured taste intensity perception to test for possible changes as a function of group during each exposure session. Participants were presented with a tray of 18 medicine cups, containing 10 ml of 3x the six taste and one mixture solution (Sucrose, Sucralose, Citric Acid, NaCl, MPG, Sucralose+Citric Acid, see section 1.3). All tastes were presented three times in a randomized order. Participants were asked to sip the solution, swirl it around in their mouth and spit it in the sink, after which they made ratings of the sweetness, saltiness, sourness, umami and general intensity of the solution. A 30 second wait-period between trials was used to rinse at least three times with deionized water. After completing the ratings for all 18 samples, participants were provided with their respective exposure beverage and were asked to finish the drink within five minutes.

*Analysis and results*

To investigate whether beverage exposure affected taste intensity ratings, we performed LMMs in R using packages LME4 and lmerTest. Models were performed for each taste solution separately. Within each model, log transformed intensity ratings were entered as dependent variable, whereas time in days, experimental group, and their interaction were entered as independent variables. Finally, participant ID was entered as random variable. To test for any interaction with ΔInsulin iAUC_0-30m_ we also performed similar models that included an interaction between change in insulin and time in days. We found no time x group interactions nor a time x group x ΔInsulin iAUC_0-30m_ interaction that survived a multiple comparison correction. The results of the intensity ratings together with the group x time interaction F-statistic derived from a mixed-effects repeated-measures ANOVA (type III) are shown in Figure S4.

**Figure S1. OGTT 120 min**

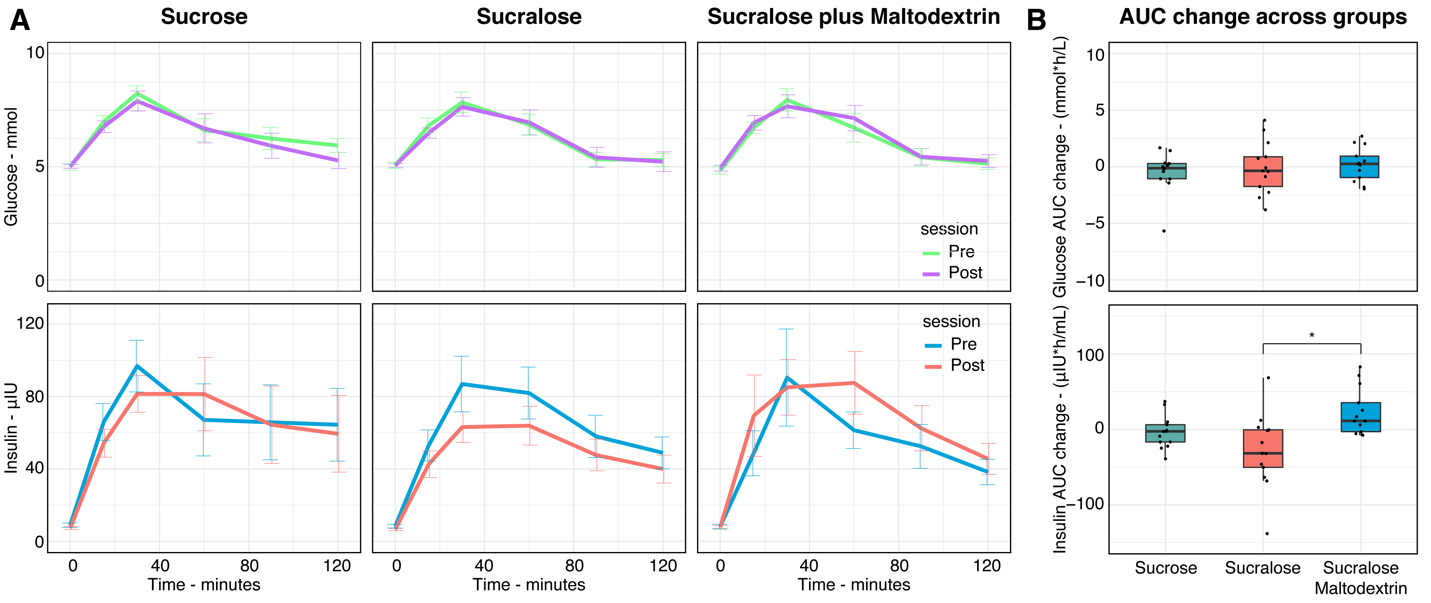

A) Full OGTT curves pre and post beverage exposure are shown for the sucrose, sucralose, and sucralose plus maltodextrin groups. The top row shows changes in blood plasma glucose while the bottom row shows changes in blood plasma insulin. B) Group differences in OGTT AUC change. Change in AUC insulin differs between the sucralose and sucralose plus maltodextrin groups (t(1,36)= 3.63, p(fdr)=0.003).

**Figure S2. Gustatory Manifold & System**

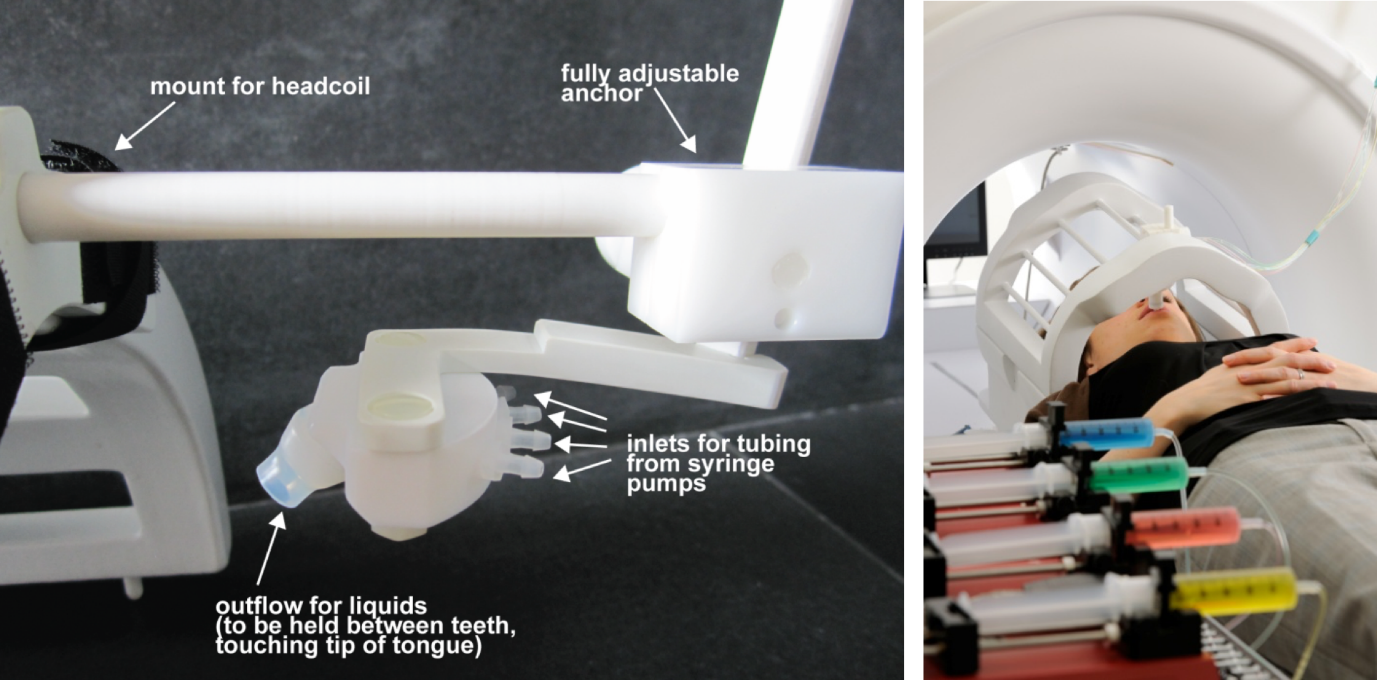

A) The gustatory manifold is photographed from a side view, showing the silicone tube that sits in between the subject’s teeth, touching the tip of the tongue. The mouthpiece has between 4-9 inlets arrayed on the semicircular bottom end parallel to the subject’s body. This configuration was designed to occupy as little space as possible inside the head coil. B) The subject is depicted in the mock scanner with gustatory manifold in position and the gustometer system in the foreground.

**Figure S3. Intensity perception changes across taste solutions and groups.**

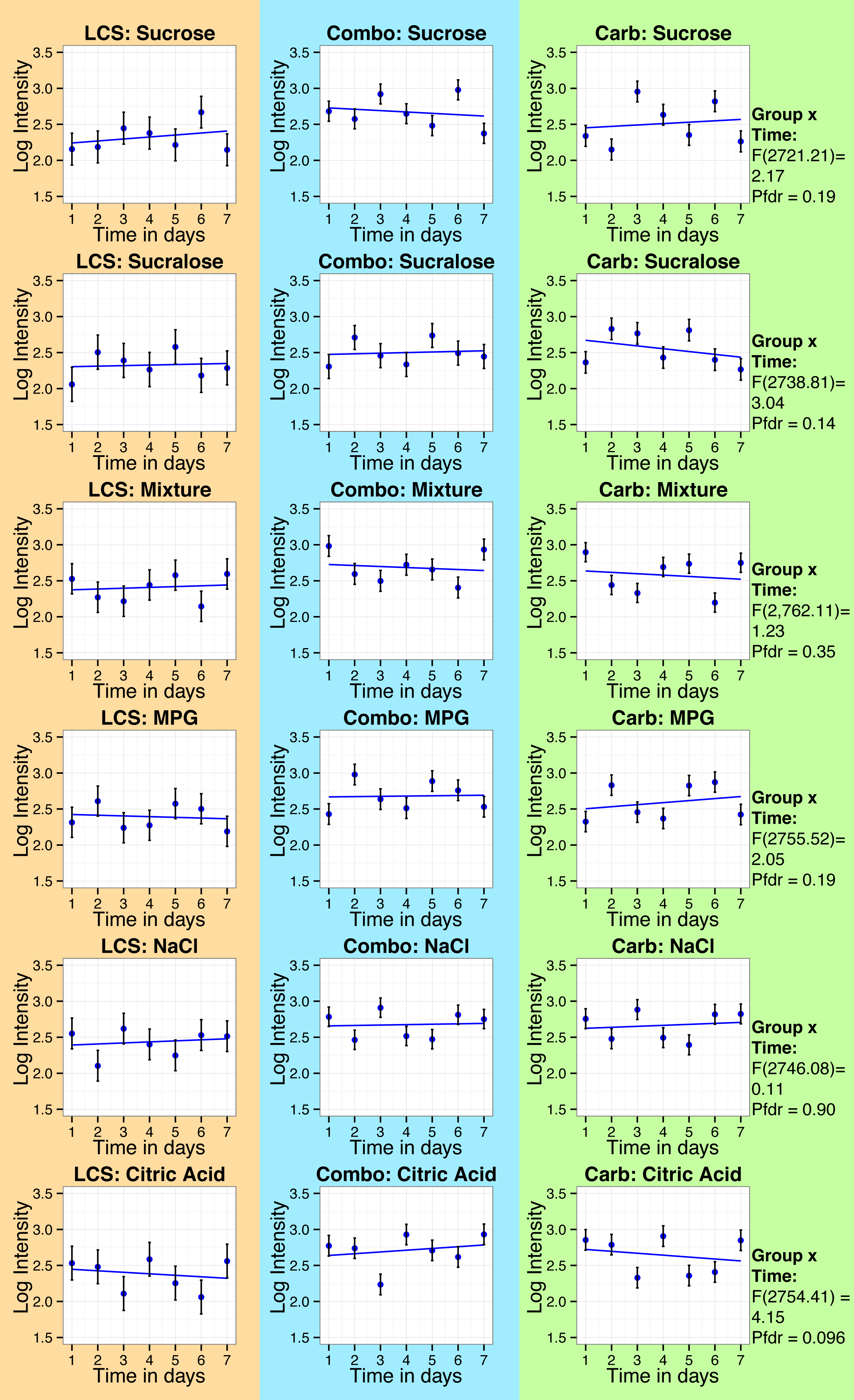

The plots indicate the average log transformed gLMS intensity ratings (±SEM) per exposure day and the linear trend across time per taste solution (rows) and group (columns). We found no group x time interaction surviving family wise error (FWE) correction. For each taste solution, F and P values are reported for the group x time interaction derived from a mixed-effects repeated-measures ANOVA.

**Figure S4. Intake, activity, fat oxidation, and energy expenditure in mice.**

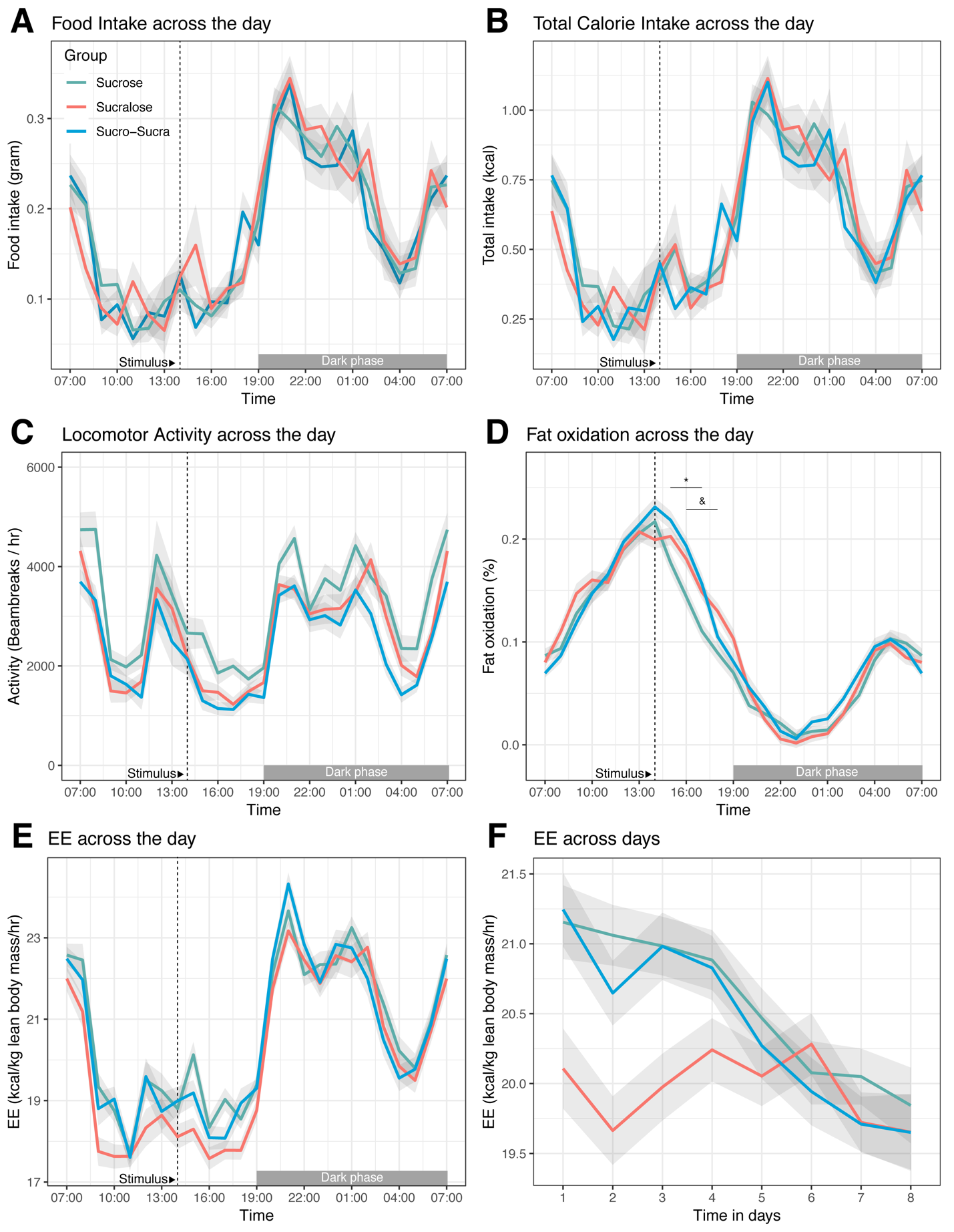

A) Hourly *ad libitum* food intake (grams). We found no differences across groups. B) Hourly total calorie intake (kcal/h). We found no differences across groups. C) Hourly activity (beambreaks/h). We found no differences across groups. D) Hourly fat oxidation (%). The sucrose group deviated from the sucralose and sucralose plus sucrose groups (false discovery rate corrected across 3x24 *t* tests; P=0.02 and P=0.007, respectively). E) Hourly energy expenditure (kcal/kg/hr). We found no differences across groups. F) Daily average energy expenditure (kcal/kg/hr). We found an interaction between group and time in days (F(2,4356.9)=8.48,p<0.001). The onset of the solution presentation is highlighted with a dashed line at 14:00h. (*) Sucrose plus sucralose vs sucrose, (#) sucrose plus sucralose vs sucralose, (&) sucrose vs sucralose. Error bands represent the standard error of the mean.

**Table S1: Detailed Participant demographics and beverage consumption**

| **Adults** | | | | |
| --- | --- | --- | --- | --- |
| Group | LCS | Combo | Carb |  |
| n | 13 | 13 | 13 | - |
| Age in years  (mean, sd) | 26.54 (3.78) | 29.15 (3.76) | 27.69 (4.19) | F(2,36)=1.46, p=0.24 |
| BMI in kg/m^2^  (mean, sd) | 22.77 (3.12) | 24.78 (2.88) | 23.62 (3.27) | F(2,36)=1.39, p=0.26 |
| Bodyfat in %  (mean, sd) | 23.6 (10.45) | 25.11 (6.40) | 29.03 (8.56) | F(2,36)=1.38, p=0.26 |
| Female:Male | 7:6 | 6:7 | 8:5 | χ^2^(2)=0.62, p=0.73 |
| Education in years (mean, sd) | 16.92 (2.10) | 17.38 (2.50) | 19.15 (3.91) | F(2,36)=2.08, p=0.14 |
| Ethnicity  Arabic  Asian  Black  Hispanic  White | 0  2  1  2  8 | 0  6  1  2  4 | 1  4  0  0  8 | - |
| Pre TLFB LCS consumption in ml over 2 weeks (median, max) | 0 (1656.12) | 0 (3193.94) | 0 (2839.06) | Kruskal-Wallis χ^2^(2)= 2.55, p=0.28 |
| Post TLFB LCS consumption in ml over 2 weeks (median, max) | 0 (4140.29) | 0 (12420.88) | 0 (3075.65) | Kruskal-Wallis χ^2^(2)= 1.48, p=0.48 |
| Pre TLFB LCS consumption in drinking days over 2 weeks (median, max)^+^ | 0 (4) | 0 (6) | 0 (8) | Kruskal-Wallis χ^2^(2)= 2.62, p=0.27 |
| Post TLFB LCS consumption in drinking days over 2 weeks (median, max) ^+^ | 0 (4) | 0 (14) | 0 (5) | Kruskal-Wallis χ^2^(2)= 1.62, p=0.45 |
| Pre TLFB SSB consumption in ml over 2 weeks (mean, sd) | 4497.45 (4168.04) | 3917.36 (3022.12) | 1574.22 (2267.77) | Kruskal-Wallis χ^2^(2)= 8.62, p= 0.01 |
| Post TLFB SSB consumption in ml over 2 weeks (mean, sd) | 4186.93 (3946.01) | 3671.67 (3240.76) | 1625.41 (1371.59) | Kruskal-Wallis χ^2^(2)= 3.82, p= 0.15 |
| Pre TLFB SSB consumption in drinking days over 2 weeks (mean, sd) ^+^ | 8.62 (5.08) | 8.08 (4.77) | 3.23 (3.77) | Kruskal-Wallis χ^2^(2)= 9.96, p= 0.007 |
| Post TLFB SSB consumption in drinking days over 2 weeks (mean, sd) ^+^ | 7.38 (4.96) | 7.85 (4.72) | 4.23 (3.54) | Kruskal-Wallis χ^2^(2)= 4.59, p= 0.10 |
| **Adolescents** | | | | |
| Group | LCS | Combo | Carb |  |
| n | 4 | 3 | 4 | - |
| Age in years | [13 14 15 17] | [16 17 17] | [14 15 15 17] | - |
| BMI in kg/m^2^ | [18.4 19.5 26.6 26.9] | [19.6 20.4 28.8] | [19 20.2 21.4 22.6] | - |
| Bodyfat in % | [14.6 36.1 36.6 28.9] | [35 24.9 18.3] | [12.5 14.0 27.9 22.3] | - |
| Female:Male | 2:2 | 3:0 | 3:1 | - |
| Education in years | [11 10 8 9] | [10 12 11] | [9 10 10 11] | - |
| Ethnicity  Asian  Black  Hispanic  White | 0  0  1  3 | 1  0  2  0 | 0  1  2  1 | - |
| Pre TLFB LCS consumption  in ml over 2 weeks | [0 0 0 0] | [0 0 0] | [0 1182.94 0]* | - |
| Post TLFB LCS consumption  in ml over 2 weeks | [0 0 0 0] | [0 0 0] | [295.74 709.76 0]* |  |
| Pre TLFB LCS consumption in drinking days over 2 weeks | [0 0 0 0] | [0 0 0] | [0 4 0]* |  |
| Post TLFB LCS consumption in # of drinking days over 2 weeks | [0 0 0 0] | [0 0 0] | [1 2 0]* |  |
| Pre TLFB SSB consumption  in ml over 2 weeks | [7275.09 709.76 2395.46 10587.32] | [4790.91 5175.37 4820.49] | [7156.79 10823.91 4820.49]* |  |
| Post TLFB SSB consumption  in ml over 2 weeks | [8546.75 0 3607.971 10528.18] | [6594.90 5175.37 8221.44] | [4199.44 11770.26 1596.97]* |  |
| Pre TLFB SSB consumption in drinking days over 2 weeks | [13 2 9 14] | [13 12 12] | [14 14 13]* |  |
| Post TLFB SSB consumption in drinking days over 2 weeks | [12 0 10 14] | [14 12 14] | [12 14 9]* |  |

*Data was not recorded for 1 participant.
^+^Total number of days on which LCS or SSB were consumed over the past 2 weeks.
Abbreviations: TLFB: Time line follow back measurement (for the preceding 2 weeks, see section 1.7); LCS: low-calorie sweetened beverage; SSB: sugar sweetened beverage.

**Table S2. Group comparisons for the OGTT indices.**

| **Measure** | **Carb**  **Mean (sd)** | **LCS**  **Mean (sd)** | **Combo**  **Mean (sd)** | **statistic** |
| --- | --- | --- | --- | --- |
| **Matsuda**  Absolute change | 0.61 (1.45) | 1.12 (2.19) | -0.35 (2.62) | F(2,36)=1.59, p=0.22 |
| Percent change | 18.41% (38.70%) | 30.73% (45.52%) | -3.02% (27.46%) | F(2,36)=2.63, p=0.09 |
| **Hepatic insulin sensitivity** Absolute change | -6.81 (16.85) | -6.15 (13.04) | 5.61 (7.04)^a,b^ | F(2,36)=3.79, p=0.03 |
| Percent change | -8.05% (81.72%) | 3.88% (110%) | 32.68% (67.84%) | F(2,36)=0.73, p=0.49 |
| **Insulin incremental AUC_0-30_**  Absolute change | -4.27 (9.71) | -4.88 (8.86) | 4.67 (5.68)^a,b^ | F(2,36)=5.43, p=0.009 |
| Percent change | -9.85% (38.93%) | -12.45% (41.59%) | 27.15% (41.00%)^a,b^ | F(2,36)=3.88, p=0.03 |
| **Glucose incremental AUC_0-30_**  Absolute change | -0.10 (0.27) | -0.10 (0.59) | -0.02 (0.29) | F(2,36)=0.19, p=0.83 |
| Percent change | -10.12 (36.38%) | 8.72% (73.88%) | 4.14% (43.75%) | F(2,36)=0.43, p=0.65 |
| **Insulin AUC_0-30_**  Absolute change | -5.06 (9.82) | -5.77 (9.91) | 4.45 (5.15) ^a,b^ | F(2,36)=5.75, p=0.007 |
| Percent change | -11.38% (30.77%) | -14.28% (33.86%) | 20.36% (29.86%) ^a,b^ | F(2,36)=4.82, p=0.014 |
| **Glucose AUC_0-30_**  Absolute change | -0.09 (0.41) | -0.11 (0.66) | 0.03 (0.32) | F(2,36)=0.33, p=0.72 |
| Percent change | -2.35% (11.79%) | -1.21% (18.07%) | 1.11% (10.48%) | F(2,36)=0.21, p=0.81 |
| **Insulin incremental AUC_0-120_**  Absolute change | -0.51 (22.33) | -24.51 (46.01) | 23.86 (32.46) ^b^ | F(2,36)=6.22, p=0.005 |
| Percent change | 5.43% (40.56%) | -14.34% (38.88%) | 24.13% (35.58%) ^b^ | F(2,36)=3.27, p=0.05 |
| **Glucose incremental AUC_0-120_**  Absolute change | -0.51 (1.24) | -0.07 (2.41) | 0.01 (1.34) | F(2,36)=0.33, p=0.72 |
| Percent change* | -18.95% (47.05%) | -45.25% (98.89%) | 7.16% (136%) | F(2,35)=0.84, p=0.44 |
| **Insulin AUC_0-120_**  Absolute change | -3.68 (21.77) | -28.14 (49.43) | 22.99 (30.92)^b^ | F(2,36)=6.58, p=0.004 |
| Percent change | 0.95% (28.18%) | -14.78% (33.82%) | 20.31% (29.04%)^b^ | F(2,36)=4.33, p=0.02 |
| **Glucose AUC_0-120_**  Absolute change | -0.47 (1.80) | -0.08 (2.32) | 0.22 (1.49) | F(2,36)=0.42, p=0.66 |
| Percent change | -3.43% (12.62%) | 0.27% (17.33%) | 2.34% (12.85%) | F(2,36)=0.53, p=0.59 |

^a^ Different from LCS, p_fdr_ < 0.05

^b^ Different from Carb, p_fdr_ < 0.05

* One extreme outlier omitted.
